# The hippocampal theta oscillation may be generated by chimera dynamics

**DOI:** 10.1101/2023.07.28.550946

**Authors:** Maria Masoliver, Jörn Davidsen, Wilten Nicola

## Abstract

The 8-12 Hz theta rhythm observed in hippocampal local field potentials of animals can be regarded as a “clock” that regulates the timing of spikes. While different interneuron sub-types synchronously phase lock to different phases for every theta cycle, the phase of pyramidal neurons’ spikes asynchronously vary in each theta cycle, depending on the animal’s position. On the other hand, pyramidal neurons tend to fire slightly faster than the theta oscillation in what is termed hippocampal phase precession. Chimera states are specific solutions to dynamical systems where synchrony and asynchrony coexist, similar to the hippocampal theta oscillation. Here, we test the hypothesis that the hippocampal theta oscillation emerges from chimera dynamics with computational modelling. We utilized multiple network topologies and sizes of Kuramoto oscillator networks that are known to collectively display chimera dynamics. We found that by changing the oscillators’ intrinsic frequency, the frequency ratio between the synchronized and unsynchronized oscillators can match the frequency ratio between the hippocampal theta oscillation (≈8 Hz) and phase precessing pyramidal neurons (≈9 Hz). The faster firing population of oscillators also displays theta-sequence-like behaviour and phase precession. Finally, we trained networks of spiking integrate-and-fire neurons to output a chimera state by using the Kuramoto-chimera system as a dynamical supervisor. We found that the firing times of subsets of individual neurons display phase precession. These results imply that the hippocampal theta oscillation may be a chimera state, further suggesting the importance of chimera states in neuroscience.

## Introduction

The hippocampus executes a complex dynamical repertoire across spatial and temporal scales to aid in behaviours that are critical for survival such as memory formation^1–10^ and navigation^10–16^. For example, the observed 8-12 Hz ‘theta’ oscillation in the local field potential organizes spikes across space^13,17–19^, time^20^, behaviour^19,21–25^, neuronal populations^26–32^, and hippocampal anatomy^33^.

Behaviourally, the theta oscillation is observed in mice and rats when they are actively engagaged in memory or navigational tasks, or during Rapid Eye Movement (REM) sleep^34–36^. The theta oscillation is critical for memory formation during these task as optogenetic or pharmacological perturbation can disrupt subsequent recall^5,6^. At the spatial level, the theta oscillation acts as a travelling wave across the septo-temporal axis of the hippocampus. Depending on the specific interneuron sub-type, interneurons primarily lock their spike times to different phases of the hippocampal theta oscillation^26–32^. Pyramidal neurons, however, fire slighly faster than the hippocampal theta oscillation, by approximately 1 Hz^5,35,36^. This frequency difference results in an effect called hippocampal phase precession, where the phase of the pyramidal neuron decreases on successive cycles.

Due to its importance in organizing hippocampal dynamics and organism behaviours across scales, the origins and mechanisms of the hippocampal theta oscillation have been intensely studied and subsequently de-bated^1–4,9,10,22–25,29,33,35–40^. The oscillation itself may be extra-hippocampal, as perturbations to the medial septum in the diagonal band of broca lead to direct changes in the hippocampal theta oscillation. Lesioning^41^, or pharmacological inhibition^6^ of the medial septum reduces the power of or eliminates the hippocampal theta oscillation while other manipulations to the medial septum can alter the theta oscillation frequency^21,42,43^. However, the whole isolated hippocampus or suitably large hippocampal slices can autonomously produce the hippocampal theta oscillation^44^. Computational modelling has shown this is possibly due to a subset of pacemaker neurons coupled with recurrent excitation, or, alternatively, as an emergent dynamical state through inhibitory neuronal interactions, or potentially emergent through local excitatory/inhibitory interactions^38–40,45–48^. Some models additionally postulate that hippocampal phase precession is inherited from other areas^49^, or created by short-term plasticity effects^50^.

In this work, rather than analyzing the network topology or biophysical mechanism of the hippocampal theta oscillation, we instead investigate the class of dynamics that can produce the observed behaviours associated with the hippocampal theta oscillation. While a straightforward oscillation as a limit cycle is one possibility, the simultaneous existence of synchronized phase-locked subpopulations of interneurons and asynchronous phase advancing pyramidal cells points to more complex dynamics. Thus, we consider chimera states, where synchronized and unsynchronized populations of oscillators co-exist^51–54^, as the dynamical state responsible for the theta oscillation’s diverse repertoire. To test the hypothesis that the hippocampus theta oscillation is a chimera state, we utilized existing computational models of chimera dynamics. The first set of models consisted of paradigmatic Kuramoto oscillators coupled with multiple network topologies that all yielded chimera dynamics^51,52,55,56^. We found that by changing the oscillators’ intrinsic oscillation frequency, the frequency ratio between the synchronized and unsynchronized oscillators can match the frequency ratio between interneurons and pyramidal neurons in the hippocampus. The unsynchronized oscillators oscillate approximately 1 Hz faster, as seen in the pyramidal neurons undergoing phase precession (see Fig. 1). These unsynchronized populations of oscillators also display sequential activity on a longer-time scale, similar to pyramidal neurons in the hippocampus during active navigation (see Fig. 1). Finally, we considered more biologically plausible models to investigate if the chimera state is responsible for hippocampal dynamics. We trained networks of spiking Izhikevich neurons^57^ to output a chimera state by using a Kuramoto-chimera system as a dynamical supervisor with FORCE training^58,59^. We found that the firing times of subsets of individual neurons display phase precession and long time scale spike sequences. These results imply that the hippocampal theta oscillation may be a chimera state, further suggesting the importance of chimera states in neuroscience.

**Figure 1.**
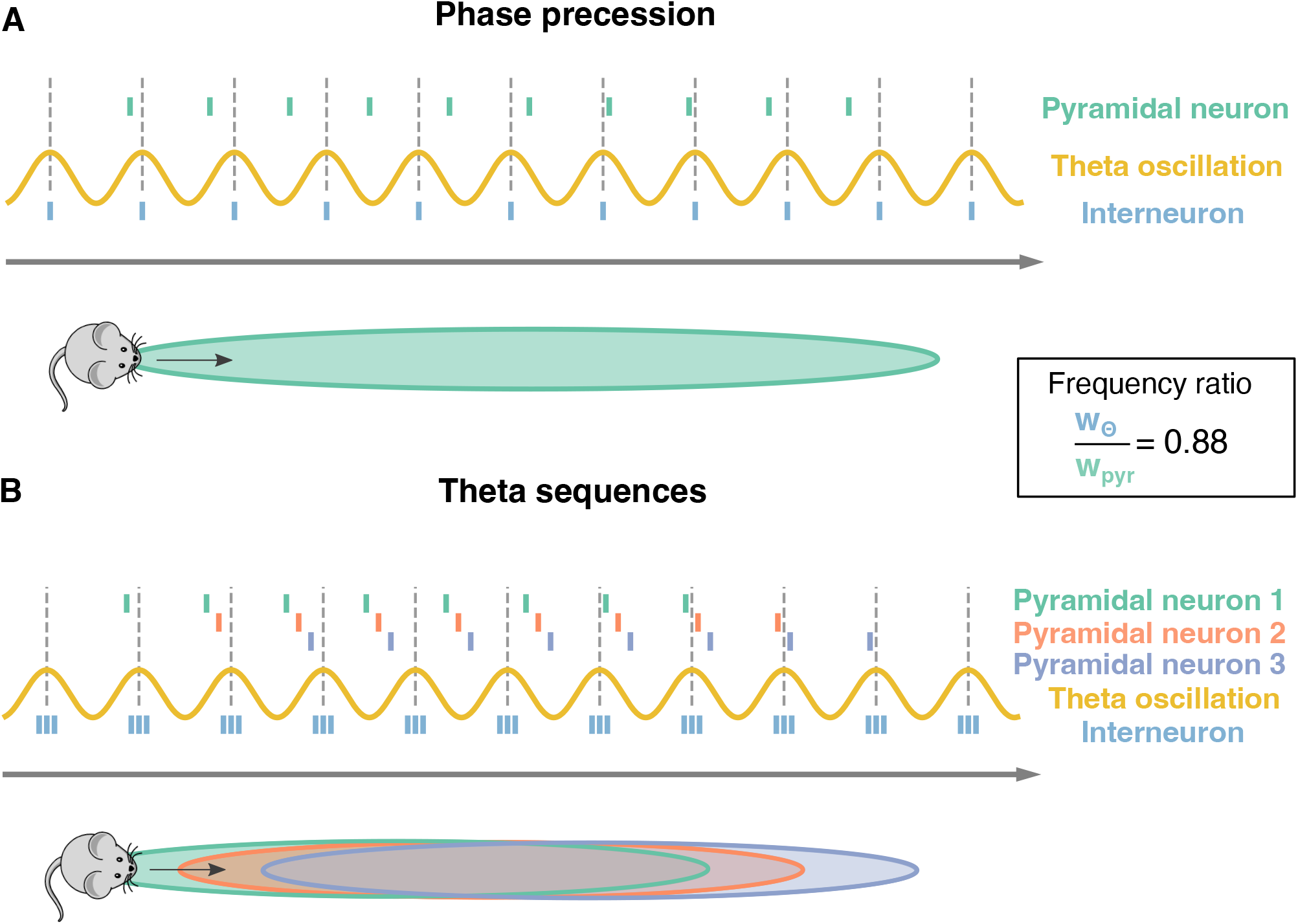
Phase precession and theta sequences. Schematic representation of phase precession and theta sequences. Dotted lines: peaks of the theta oscillation (yellow sinusoidal curve). The distance between two subsequent lines defines one theta cycle. **A** As a mouse moves along a track (grey arrow); a pyramidal neuron starts firing as the animal enters the pyramidal neurons’ place field (green surface). While the actions potentials from the pyramidal neuron (green ticks) happen earlier at each theta cycle, the ones from an interneuron (blue ticks) are usually synchronized to the theta oscillation. This phase advancement from the theta cycle is known as phase precession. The ratio between theta oscillation’s frequency and pyramidal neuron’s frequency is approximately 0.88 = 8 Hz/9 Hz as pyramidal cells tend to fire at approximately 1 Hz faster than the 8 Hz theta oscillation **B**. Three pyramidal neurons undergoing phase precession and their corresponding place fields are considered (green, red and purple). Having multiple neurons leads to sequences of spikes within a theta oscillation, known as theta sequences. Three interneurons (blue ticks) are depicted as well.

## Results

### Analyzing Existing Chimera-Inducing Network Topologies

To investigate if chimera dynamics are a potential mechanism for the neuronal dynamics associated with the hippocampal theta oscillation, chimera dynamics were first simulated in pre-existing models to test the hypothesis that parameter ranges that exhibit hippocampal-like dynamics (i.e. phase precession and sequential content) could readily be determined.

Two paradigmatic model versions each generating a different chimera state — a chimera on a ring and a two-population chimera, respectively — were considered. The chimera on a ring arises for *N* = 500 nonlocally coupled identical Kuramoto oscillators (see Fig. 2A and Methods, Eq. (2)), whereas the two-population chimera arises for two weakly coupled populations, each one formed by 3 globally coupled Kuramoto oscillators (see Fig. 2E, Supplementary Figure 1A and Methods, Eqs. (3, 4)). Depending on the parameter values and on the initial conditions, both network topologies can display different dynamics: Either a fully synchronized state where all oscillators are in phase or chimera states where one sub-population of neurons is synchronized while the other sub-population oscillates asynchronously (see Video 1, supplementary material).

**Figure 2.**
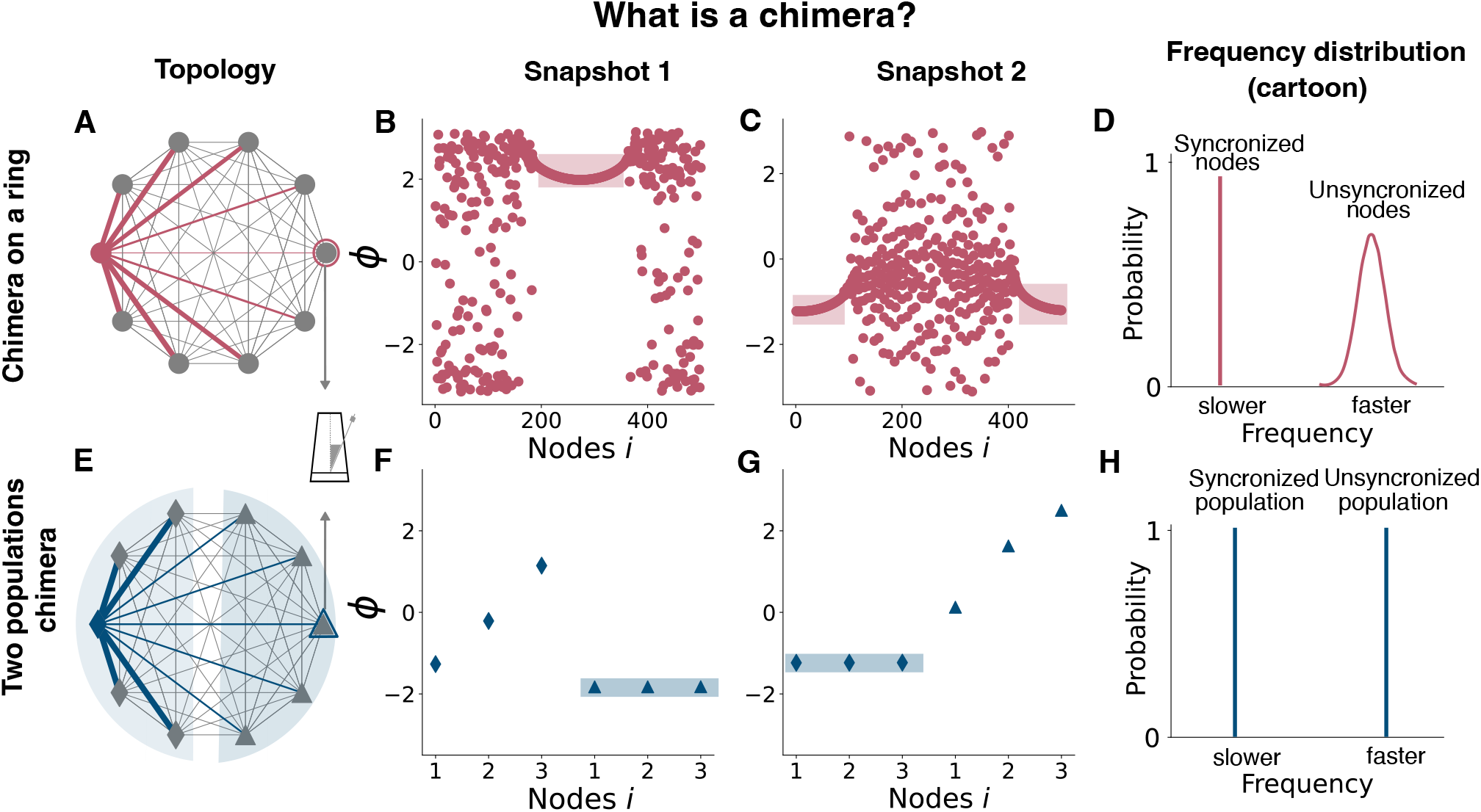
Chimera states in two networks of Kuramoto Oscillators. Schematic representation of two different chimera states: the chimera on a ring (top) and the two-population chimera. For both topologies, each node is a Kuramoto Oscillator. **A** Diagram showing the coupling scheme needed to observe a chimera on a ring. A nonlocal coupling rule is used, see Eq. (2) for details. For clarity, only the coupling for a single node or oscillator (pink node) has been depicted (pink edges). Edge thickness represents the weights of a connection. **B** Snapshot of oscillators’ phases at a given time. Light pink rectangle denotes the synchronized nodes. **C** Snapshot of oscillators’ phases at a different time. Light pink rectangle denotes the synchronized nodes. **D** Schematic representation of the ring oscillators’ frequency distribution. The synchronized nodes oscillate at the same frequency (within them) but at a slower pace than the unsynchronized ones. **E** Diagram showing the coupling scheme needed to observe a chimera on two populations: two populations (diamonds and triangles) are weakly coupled between each other and strongly coupled within, see equations (3) and (4) for details. For clarity, just the coupling for one node (blue) has been depicted (blue). Edge thickness represents the weights of its connection. **F** Snapshot of oscillators’ phases at a given time. Light blue rectangle marks the synchronized population. **G** Snapshot of oscillators’ phases using different initial conditions or after the system is externally perturbed (see Supplementary Figure 2). Light blue rectangle marks the synchronized population. **H** Schematic representation of the ring oscillators’ frequency distribution. The synchronized population oscillates at a slower pace than the unsynchronized one. The nodes for each population oscillate at the same frequency.

First, the model parameters of the dynamical equations (Eq. (2) and Eqs. (3, 4)), respectively) were set to well known or classical parameter regimes where chimera dynamics readily emerge^56,60^. The parameters *A* and *β* affect the coupling strength and the phase difference, respectively, and take different values for the two different systems, see Table 1 for details. For the chimera on a ring, the synchronous subpopulation of oscillators is non-static, and drifts slowly around the ring. Oscillators drift in and out of the synchronous sub-population, while the others oscillate asynchronously (see Video 2, Fig. 2B and Fig. 2C). In contrast, for the two-population chimera, the chimera state is static: one population fully synchronizes (triangles in Fig. 2F) while the other one does not (diamonds in Fig. 2F). Unless the system is perturbed, the synchronized and unsynchronized populations remain fixed. The identity of the synchronized or unsynchronized population depends on the initial conditions (Fig. 2F,G). The synchrony profile between the two populations can be exchanged by externally perturbing the system, where the synchronized and unsynchronized populations swap. For example, in Supplementary Figure 2, the triangle population is synchronised before a perturbation, and after a perturbation, the oscillators move to an asynchronous regime (vice versa for the diamond population).

**Table 1.** Parameters for the model chimera-on-a-ring, described in Eq. (2) and for the two populations chimera, described in equations (3) and (4).

| **Chimera on a Ring** |  |
| --- | --- |
| Parameter | Value |
| A | 0.95 |
| $\beta$ | 0.2 |
| N | 500 |

| **Two populations Chimera** |  |
| --- | --- |
| Parameter | Value |
| A | 0.1 |
| $\beta$ | 0.025 |
| n | 3 |
| $\tau$ | 1/0.012 |
| $\mu$ | 0.18 |
| $\nu$ | 0.15 |

For the chimera on a ring, the synchronized and unsychronized populations drift (Video 1, Video 2 and Fig. 2B,C). An external perturbation, in this case, is not necessary to change the oscillators’ synchrony profile. The identity of the neurons that constitute the synchronized population slowly drifts around the ring as a slowly moving travelling wave. As the drift’s period is much larger than the oscillations’ period, we can study the differences between the two domains, synchronized and unsynchronized (see ref.^55^ for details on the drift).

### From a chimera state to hippocampal phase precession

With the classical chimera dynamics reproduced, we investigated how to explicitly draw a mapping between the Kuramoto networks and hippocampal dynamics. Each neuron has more complex dynamics than those of a Kuramoto oscillator. Specifically, neurons emit spikes when their inputs are sufficient to reach a threshold. Thus, each oscillator’s continuous time-series was converted into spike trains via a Poincare Map. Each time any Kuramoto oscillator’s phase reaches 2*π*, a “spike” is generated at the time that this occurred (*θ*_*j*_(*t*^*^) = 2*π*) as depicted in Fig. 3C (see Methods for details). With a spike-generating Poincare map, the “spikes” generated by the chimera on a ring (Fig. 3D) and for the two-population chimera (Supplementary Fig 1C) can be analyzed.

**Figure 3.**
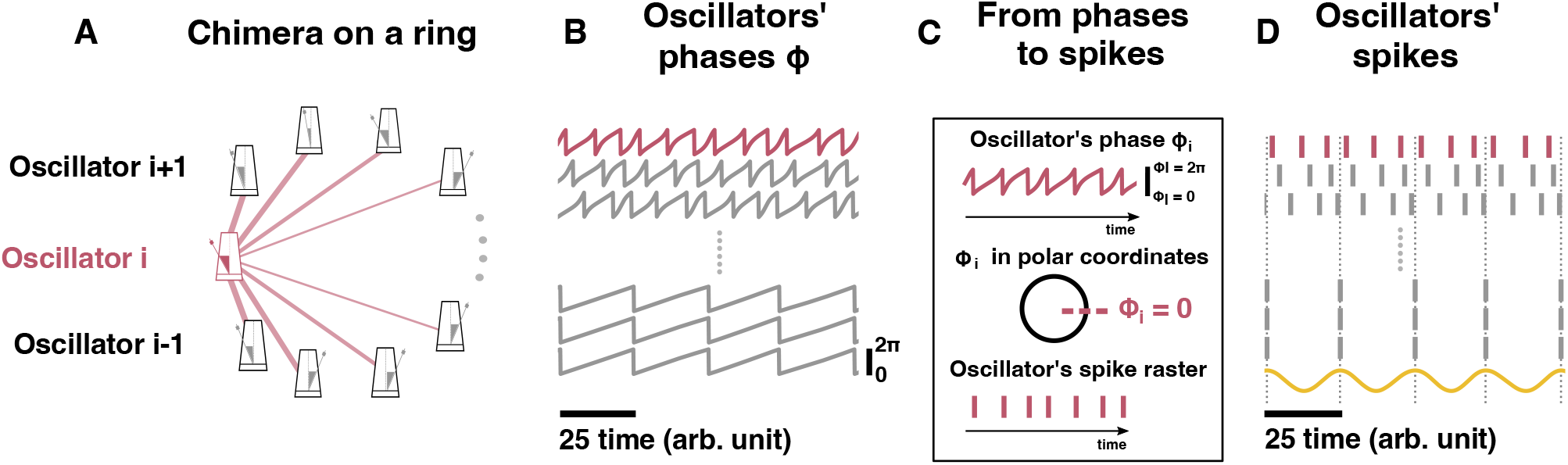
From a chimera state to hippocampal phase precession. **A** Schematic representation of the chimera state on a ring (see Eq. (2) for details). For clarity, the coupling for oscillator *i* (pink or dark grey metronome for b/w printing) has been depicted (pink or dark grey edges for b/w printing). Edge thickness represents the weights of connections. **B** Time-series of different oscillators: some are asynchronous (top) and some are synchronous (bottom). **C** Cartoon explaining the transformation from the oscillators’ phases to a putative “spike”: every time ***ϕ***_*i*_ = 0, a spike occurs. **D** Oscillators’ spike raster plot: panel **B** transformed into a raster plot. The sinusoidal curve (yellow) represents the macroscopic theta oscillation observed in an LFP. The theta oscillation is computed as the mean of cos(***ϕ***_*j*_), where ***ϕ***_*j*_ corresponds to the phase from the synchronized oscillators. Dotted lines correspond to the peaks of the sinusoidal signal and to the spikes of the synchronized nodes. Model parameters: *A* = 0.95, *β* = 0.2 and *N* = 500, *ρ* = 1.

In order to measure phase-precession, an equivalent component to the hippocampal local field potential in the Kuramoto network is required. The hippocampal LFP is a macroscopic observable that represents the bulk-flow of ions across the membranes of neurons near the recording electrode. During *in vivo* recordings, the hippocampal LFP is typically converted into a phase (for example with a Hilbert transform). Interneuron’s and sometimes pyramidal neurons lock to phases of the hippocampal LFP, while other pyramidal neurons fire at a slightly faster rate.

Given the locking of synchronized sub-populations to the hippocampal LFP, a phenomenological LFP was computed as follows: the cosine of the phase of each oscillator in the synchronized population was computed (cos ***ϕ***_*j*_) and globally averaged over the synchronized population. The LFP can also be computed as the mean over all oscillators (both synchronized and unsynchronized): 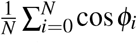(Supplementary Fig. 3A and 3B).

Interestingly, we observed phase advancement from the unsynchronized oscillators when compared to the synchronized ones (Fig. 3D, and supplementary figure 1C). While this is similar in principle to phase precession, where the unsynchronized pyramidal neurons fire slightly faster than the local-field-potential, the frequency ratio between the synchronized oscillators and the unsynchronized is different from those observed experimentally. For every synchronized spike we get approximately three unsynchronized ones, which roughly gives us a ratio of ≈0.33 (Figure 3D). In the hippocampus, pyramidal neurons fire at approximately 9 Hz, while the theta oscillation observed in the LFP is approximately 8 Hz, which yields a a ratio of ≈ 0.88.

### Changing the chimera state by changing the intrinsic frequency

Next, we investigated if the parameters in both models could be varied to both preserve the chimera state, and obtain a frequency ratio closer to that of hippocampal phase precession (≈0.88). Accomplishing this in both models would indicate that one can generically obtain hippocampal-like dynamics in chimera systems. To start, the intrinsic frequency parameter (*ρ*) was varied in Eqs. (2), (3) and (4). This acts as the fundamental driving force for an oscillator and causes the oscillator to intrinsically oscillate when no coupling is present. Thus, it is directly comparable to the applied current *I* typically considered in neuron models as higher applied currents lead to faster firing rates (more spikes per unit time).

As *ρ* was varied, the oscillating frequency for each oscillator was quantified as follows: the mean phase velocity Ω_*i*_ was computed for oscillator *i* to determine its frequency. As the driving frequency *ρ* interacts with the coupling in a non-trivial way, the frequencies must be computed numerically. For a given oscillator *i* and a given amount of time Δ*t*, the number of rotations around the origin (or equivalently, the number of spikes fired) was summed and multiplied by 2*π* (see methods and Eq. (5) for details). This was was then divided by Δ*t* to yield the rotations.

To see if the chimera dynamics could mimic hippocampal observations, we focused on the mean phase velocity ratio ⟨Ω _*ratio*_⟩. This ratio was computed as the average of the mean phase velocity ratio between the synchronized and unsynchronized populations as a function of *ρ*, as shown below (see Methods for details):

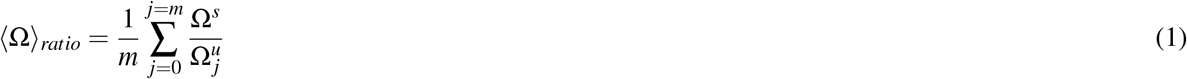

As the intrinsic oscillation frequency (*ρ*) increases, the oscillation frequency of both the synchronized and unsychronized oscillators in the coupled network increases, but the frequency difference between the synchronized and unsynchronized domains decreases. This was quantified for *ρ* = 1.8 (Fig. 4C) and *ρ* = 2.8 (Fig. 4D) and, more generally, for the mean phase velocity ratio as a function of *ρ* (Fig. 4E). As *ρ* was varied, **Ω**^*u*^ varied over a range which was bounded by a minimum Ω_*min*_ and a maximum value Ω_*max*_. For *ρ* = 1.8, (Ω_*min*_, Ω_*max*_) = (1.056, 1.565) while for *ρ* = 2.8, they increase to (Ω_*min*_, Ω_*max*_) = (2.055, 2.545) and we achieve ⟨Ω⟩ _*ratio*_ ≈0.88 for that value (Fig. 4E). As *ρ* is increased further past this value, the ratio slowly increases until the chimera state collapses and all oscillators synchronize (see supplementary figure 4).

**Figure 4.**
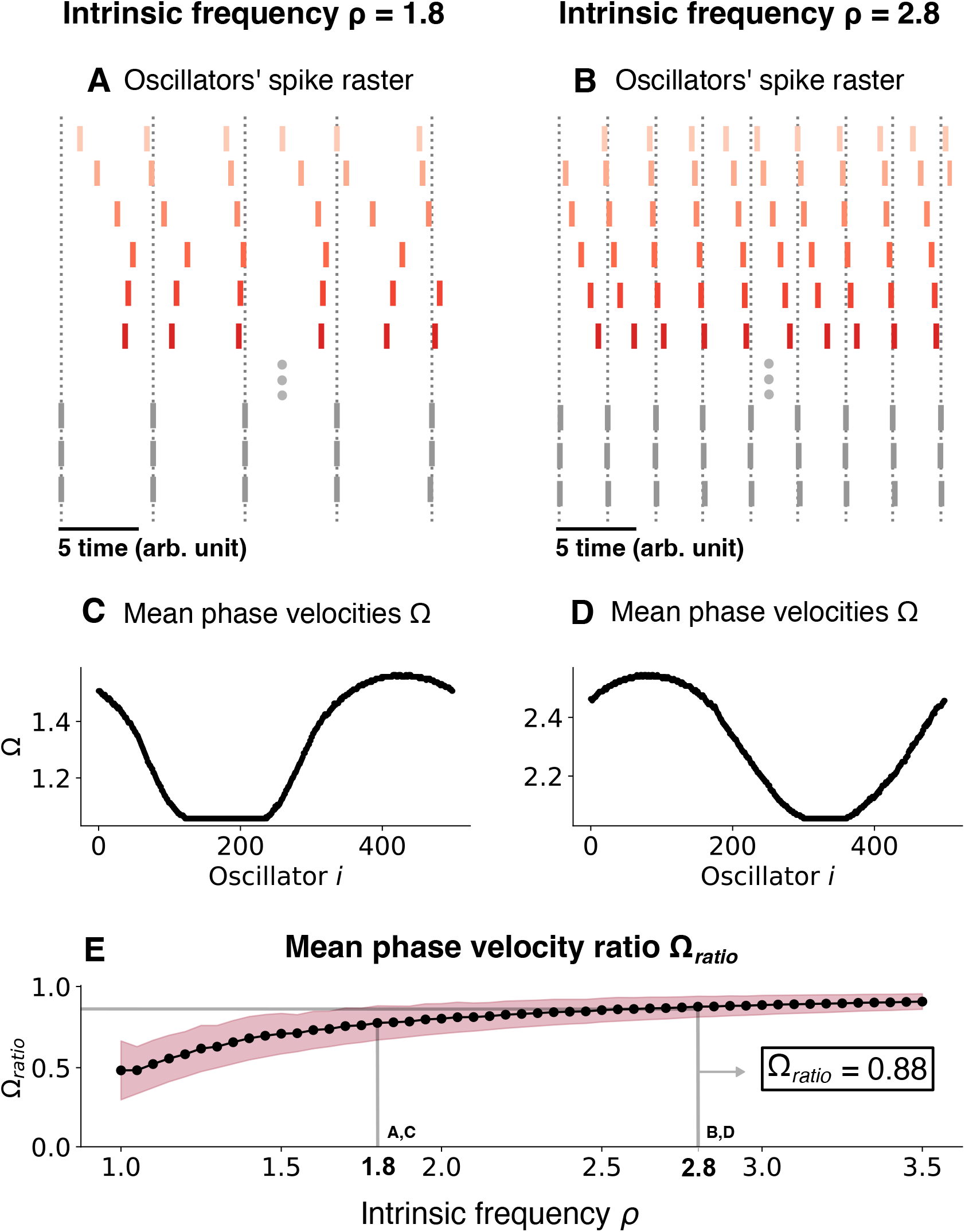
Changing the chimera state by changing the intrinsic frequency. **A** Oscillators’ spike raster plots for *ρ* = 1.8 and **B** *ρ* = 2.8, respectively. Note the theta sequences contained within a single oscillation cycle. Dotted lines correspond to the spikes of the synchronized nodes (grey ticks). **C** Mean phase velocity profile for *ρ* = 1.8 and **D** *ρ* = 2.8, respectively. **E** Mean phase velocity ratio ⟨Ω⟩_*ratio*_ as a function of the intrinsic frequency *ρ*. The pink region indicates the spread of the mean phase velocity ratio as computed by the standard deviation of ⟨Ω⟩_*ratio*_.

Next, we tested if this was a generic response by considering the two-population chimera model (see Supplementary Figure 5). Once again, we found that the ⟨Ω _*ratio*_⟩≈0.88 can occur for a specific *ρ* due to the slow gradual increase in ⟨Ω _*ratio*_⟩as a function of *ρ*. Thus, the phase precession regime of classical chimera models is seemingly robust and generic.

Finally, we investigated what the net impact of the coupling was. That is, we considered how the mean phase velocity for both domains (synchronized and unsynchronized) and for both network topologies compares to the mean phase velocity of an uncoupled oscillator. In the latter case, the mean-phase velocity is given by

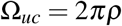

Interestingly, regardless of the network topology, the net effect of the coupling was always inhibitory: The oscillators fire at a faster frequency when uncoupled, rather than when coupled into a chimera state in both network topologies (see Supplementary Figure 6).

### Phase precession in a chimera-trained spiking neural network

Chimera dynamics in networks of Kuramoto oscillators with different network topologies can be altered by changing one parameter, the intrinsic driving frequency, to mimic a hippocampal-like phase precession regime. Despite the general nature of these results, the Kuramoto-oscillator network is phenomenologically different from the neurons and synaptic connections in the hippocampus. Thus, we sought to determine if embedding a chimera state into a spiking-neural-network would still yield hippocampal phase precession, and a global theta-oscillation.

A chimera state can be “embedded” in a recurrent neural network by training the network to output a chimera, as seen in^61^. To test if such an embedding was applicable in a spiking network, we trained a spiking neural network using the FORCE method^58,59^ to output the two-population chimera, described by Eqs. (3) and (4). This network was constrained with Dale’s law, with a proportion of the neurons being excitatory, and the rest inhibitory. Initially, the individual neurons (modeled using the Izhikevich model, see Methods for details) are sparsely connected (to support the learning process^58^) with a set of static weights *G****ω***_0_ which initiate the neurons’ rate ***r***(*t*) into a high-dimensional chaotic regime. During the training period a second set of weights *Q****ηd***^*T*^ is added to *G****ω*** _**0**_ and changes the connections between neurons such that the network’s output (defined as ***d***^*T*^ ***r***) equals the desired dynamics. The desired dynamics or supervisor are cosines of the phases of a two-population Kuramoto oscillator network in the chimera regime. At each time step, ***d*** is updated using the Recursive Least Squares (RLS), which minimizes the sum-squared difference between the network output and the two-population chimera. The network has learned when for a fixed value of ***d*** it is able to mimic the desired chimera dynamics (see Supplementary Figure 7). A specific example is shown in Fig. 5.

**Figure 5.**
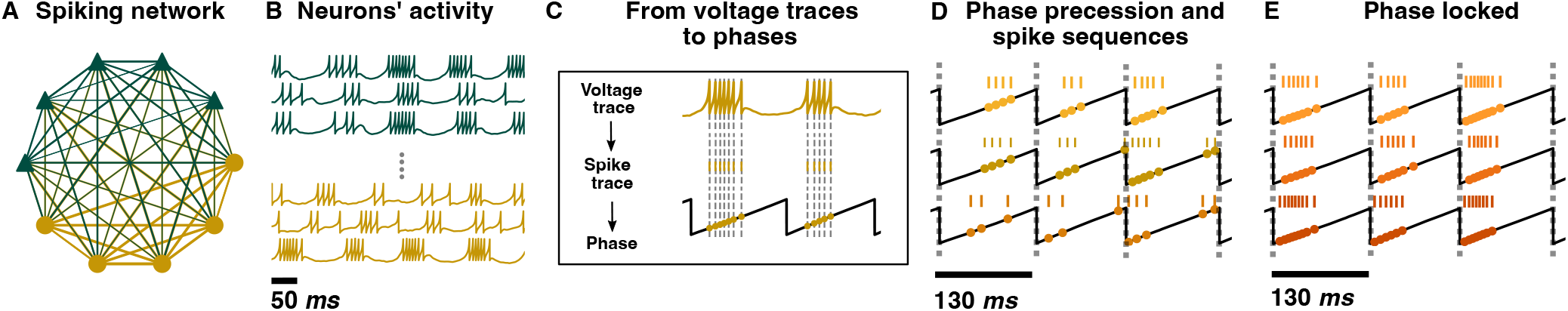
Phase precession in a chimera-trained spiking neural network. **A** A cartoon schematic of the spiking neural network with Dale’s law. Each node represents either an excitatory (green or dark grey triangle for b/w printing) or inhibitory (yellow or light grey circle for b/w printing) neuron. The network respects Dale’s law: an excitatory (inhibitory) neuron will only excite (inhibit) its connections, regardless of the neuron target type. As a result, excitatory (inhibitory) neurons only have green or dark grey for b/w printing (yellow or light grey for b/w printing) outgoing connections. Edge thickness represents the weight of each connection. **B** Voltage traces for excitatory (green or dark grey triangle for b/w printing) and inhibitory (yellow or light grey round for b/w printing) neurons. **C** From top to bottom: the voltage trace of an inhibitory neuron from the spiking neural network, its correspondent spike sequence and its projection to the phase 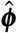 of the synchronized population (black trace), which represents the theta oscillation. Grey dotted lines mark every time the voltage reaches its peak *v* = 30 mV and a spike is generated. **D** Example of phase precession and spike sequences from three inhibitory neurons. For each neuron, the spike sequence and its projection into the phase 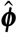 is plotted. Grey dotted lines mark every time 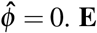. **E** Example of three phase locked inhibitory neurons. Grey dotted lines mark every time 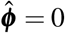.

In order to assess if the individual neurons of the spiking network show phase precession, the voltage traces were transformed into phases (Fig. 5C). The spike times were transformed into phases by using a linear interpolation to approximate the phase at each spike time with the phase of one of the synchronized components of the network output 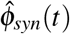 We found that phase precession occurred generically for many of the neurons sampled, where neurons displayed decreasing burst phases based on subsequent cycles (Fig. 5D). However, some of the neurons were primarily phase locked (Fig. 5E).

## Discussion & Conclusions

Since their discovery, chimeras have been extensively modeled, applied, and recently experimentally realized in the study of complex oscillatory systems^53,54,62–65^. More recently, attempts have been made to link them directly to brain dynamics, using largely modelling studies and different coupling topologies^66^. This includes chimeras in oscillating brain networks^67,68^, three-dimensional chimeras in spiking neuronal networks^69^, chimeras in heterogeneous networks^56,70^ and the robust emergence of chimeras in recurrent neural networks^61^ as well as limited experimental studies^71^. Yet, their potential functional role in brain dynamics has remained largely elusive. Chimera’s have been hypothesized to be the dynamical state dolphins, birds, and other animals that need to navigate over large ranges in 3-dimensions utilize to sleep, where half the brain is in a synchronized sleep state while the other half is in an asynchronous awake state^54^. Similarly, chimera’s might potentially play a role in memory consolidation related to REM and non-REM sleep^72^.

Here, we utilized computational modelling to test the hypothesis that the hippocampal theta oscillation may be a chimera state. By modifying the intrinsic frequency parameter in the Kuramoto oscillators exhibiting a classical chimera, and using a spike-generating Poincare map, we found that chimera dynamics readily produced theta-phase precession-like observations over a range of values. The oscillators in the asynchronous group fired slightly faster (∼1 Hz) than those in the synchronous group, resulting in theta phase precession. The spikes generated by these oscillators also displayed theta-sequence-like activity We found that the net coupling in both the chimera-on-a-ring and two-population chimera was inhibitory, as deactivating the coupling resulted in a higher mean-phase-velocity than with the coupling in place. Finally, we embedded a chimera state into a spiking neural network of Izhikevich neurons with Dale’s Law constraining the connection weights. Despite the embedded nature of the chimera, at the micro-scale, the spiking neurons still displayed phase-precession (asynchrony) and phase locking (synchrony), the observable features of the chimera. This is despite the heterogeneity in the coupling the neuron’s display. Collectively, this study is the first to postulate and test the hypothesis that the hippocampal theta oscillation is a chimera state.

Interestingly we found that the synchronized and unsynchronized population(s) can drift, and thus the designation as being part of the synchronized and unsynchronized population is non-static while the global chimera state persists. This feature is generic to many chimera models, especially in chimera models involving 3-dimensional structures^63,73–75^. This is consistent with the fact that phase-precessing pyramidal neurons are not fixed and change their dynamics over time^76^. This distinguishes the chimera hypothesis from other hypotheses for the generation of hippocampal phase precession where the phase-precession effect is in some cases fixed by either the local or global connectivity (e.g.^38,46,48^). Indeed, this is the intrinsic difference between chimera dynamics, and other models of phase precession: chimera’s allow considerable flexibility in which neurons are phase precessing dependent on changing the initial conditions or external inputs or perturbations.

Chimera states have proven to be ubiquitous and robust in nature, whether implemented as collections of simple pendulums or metronomes, or in the underlying dynamics behind chemical reaction equations. However, the heterogeneity and noise present in biological systems may destabilize these dynamical states. Here, we show that Chimera states are intrinsically linked with hippocampal phase-precession, and possibly present the first biological chimera state observable at a cellular level.

## Methods

### Chimera on a ring

The chimera state on a ring was obtained from the integration of *N* Kuramoto oscillators with nonlocal coupling. The equations are given by^55^:

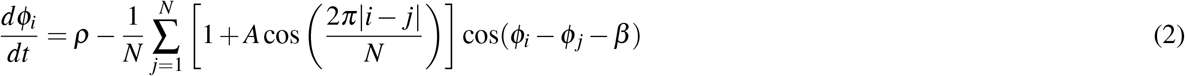

with *i* = 1, …, *N*, where ***ϕ***_*i*_ is the oscillator’s phase and *ρ* is the oscillators intrinsic frequency. To obtain a stable chimera we integrated Eq. (2) using chimera-like initial conditions and we set *A* = 0.95, *β* = 0.2 and *N* = 500 as in ref.^56^. We chose *A* and *β* from the (*A, α*) parameter plane in which the chimera state exists (see ref.^55^ for details) and set *N* large enough (*N >* 50) such that for this type of network the chimera did not collapse (note that the number of oscillators can be reduced when using a different topology as the one used in Eq. (2), see Supplementary Material and ref.^60^ for examples). We obtained chimera-like initial conditions by randomly selecting the same phase for half of the network. The phases for the other half were selected from a uniform distribution between [0, 2*π*]. See ref.^61^ for more details and refer to Supplementary Material for the exact initial conditions used in this publication. The equations were integrated using the Euler method with an integration step of *dt* = 10^−3^. Note that all equations are dimensionless, however time can be rescaled so that a single unit of time, which corresponds to 8 cycles of the synchronized population, can be rescaled to 1 second which yields an 8 Hz (theta) oscillation.

### Two-population chimera

The two-population chimera consists of two populations of *n* Kuramoto oscillators each. The phases of the oscillators for group 1 and group 2 are given by 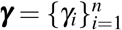and 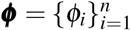, which are governed by the following equations:

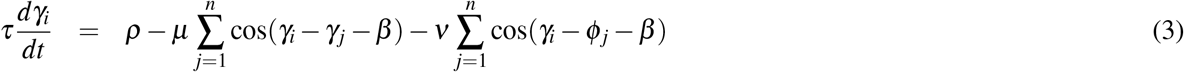

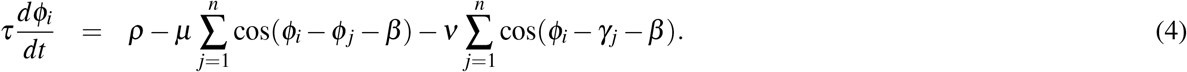

The coupling within groups is given by 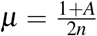 and between groups by 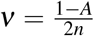, with *ν < μ*, 0 ⩽ *A* ⩽ 1 and *n* = 3. To obtain a stable chimera, we simulated equations (3 and (4 with appropriate initial conditions as in ref.^61^ and we fixed *β* = 0.025 and *A* = 0.1. The temporal component 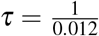 is used to slow down the chimera dynamics, which is needed in order to successfully train the spiking recurrent network. Is for that value that we get 8 and 9 oscillations, for the synchronized and the unsynchronized populations, respectively, in 1 second. Figure 2E and Supplementary Figure 1A illustrate the network architecture, supplementary Figures 5A and 5B depict the chimera dynamics. By tuning the parameter *ω* we obtain the desired ratio between the mean phase velocity of the synchronized and unsynchronized populations as shown in Supplementary Figures 5E, 5F and 5G. The equations were integrated using the Euler method with an integration step of *dt* = 10^−3^. Note that all equations are dimensionless, but can be rescaled in time as described in the chimera-on-a-ring case.

### From phases to spikes

Since the oscillators are periodic and oscillate between 0 and 2*π*, we can transform the oscillators’ time-series (Fig. 2B and Supplementary Fig. 1A) into a raster plot. Every time the oscillator’s phase ***ϕ***_*i*_ = 0 a spike is drawn (Fig. 3C). The resulting raster plot from the aforementioned time-series is shown in Fig. 3D for the chimera on a ring and in Supplementary Figure 1C for the two populations chimera.

### Mean phase velocity

The mean phase velocity^77^ for a given oscillator with phase *θ*_*i*_ is defined as :

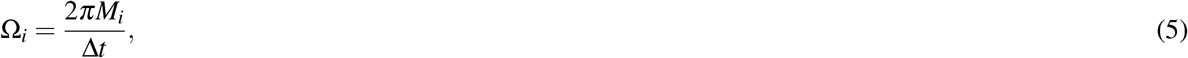

where *M*_*i*_ is the number of complete rotations around the origin performed by the *i*th oscillator during the time interval Δ*t* = 1000. It acts as a measure of the oscillating frequency for each oscillator. Given a ring of *N* oscillators, it is denoted as **Ω** = Ω_*i*_, with *i* = 1, 2, … *N*. Having different mean phase velocities for the synchronized and unsynchronized domain is typical for chimera states. In particular, for ring-like topologies it is common to have an arc-like profile of mean phase velocities for the unsynchronized domain, denoted as 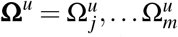 (where *u* stands for unsynchronized and *m* is the total number of unsynchronized oscillators) and equal mean phase velocities for the synchronized domain^62,64,78^ given by 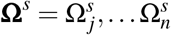 (where *s* stands for synchronized and *n* is the total number of synchronized oscillators). Since the oscillators are synchronized they have the same mean phase velocity Ω*s*, therefore we can simplify **Ω**^*s*^ to a unique value given by Ω_*s*_. We will use Ω_*s*_ to identify which oscillators synchronize and which do not, given that the synchronized domain oscillates at a slower pace than for the unsynchronized one Ω*s <* **Ω**^*u*^∀ *j* ∈ *m*. In order to identify Ω*s*, one can simply compute the minimum of **Ω**.

For the two-population chimera, both domains (synchronized and unsynchronized) have equal mean phase velocities (different between domains but equal within). Also for that topology, the synchronized population oscillates at a slower pace than for the unsynchronized one: Ω*s <* Ω*u*.

### Mean phase velocity ratio

#### Chimera on a ring

For each intrinsic frequency *ρ* we compute the mean phase velocity ratio, which measures the relation between the mean phase velocity of the synchronized domain versus the unsynchronized one. We define the mean velocity ratio as follows:

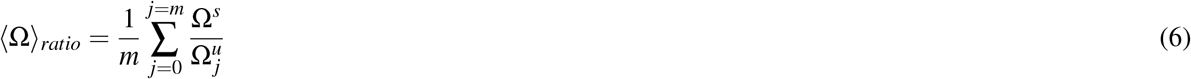

As with any average, there is a standard deviation associated with it:

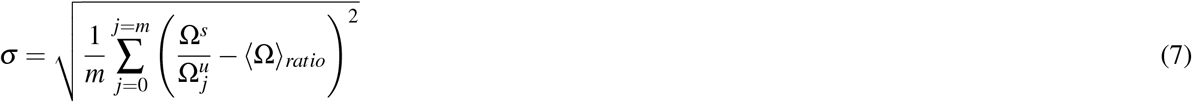

#### Two-population chimera

For the two-population chimera, the mean phase velocity is simplified, since we do not have a unique value only for Ω^*s*^ but also for Ω^*u*^. We can rewrite Eq. (7) as:

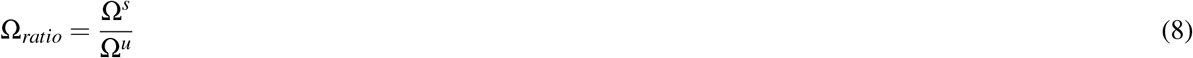

since we do not have a set of values for Ω_*u*_ (see Supplementary Figures 5E and 5F), there is no variation when computing Ω_*ratio*_ as depicted in Supplementary Figure 5G.

### Spiking neural network equations and the FORCE method

The spiking neural network consists of coupled Izhikevich neurons^57^, with their dynamics given by the following equations:

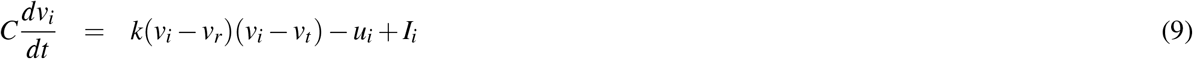

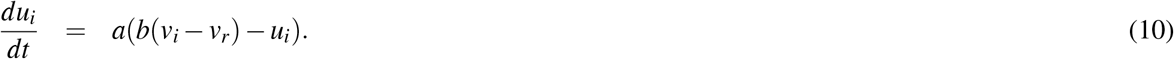

The quantity *v*_*i*_ is the voltage variable. Neuron *i* fires a spike when *v*_*i*_ reaches a voltage peak *v*_*peak*_ and it is instantly reset to a potential *v*_*reset*_. The adaptation current is given by *u*_*i*_, which increases an amount *d*_*u*_ every time a spike is fired and which in turn slows down the production of spikes. The current *I*_*i*_ is given by *I*_*i*_ = *I*_*bias*_ + *s*_*i*_, where *I*_*bias*_ is a fixed value and *s*_*i*_ are the synaptic currents for neuron *i*, given by

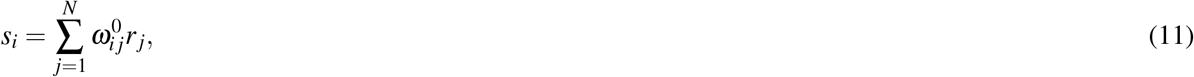

where *N* is the total number of neurons. The matrix 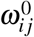controls the magnitude of the postsynaptic currents arriving at neuron *j* from neuron *i*. The parameter *C* represents the membrane capacitance, the parameters *v*_*r*_ and *v*_*t*_ denote the resting and the threshold membrane potential, respectively. The parameter *a* is an equivalent of the time constant for the adaptation current *u*_*i*_. The parameter *b* controls the resonance properties of the model and *k* controls the half-width of the action potentials. The numeric parameters of the model are listed in table 2, we used the same parameters as in^59^. The spikes are filtered with a double exponential synapse, given by:

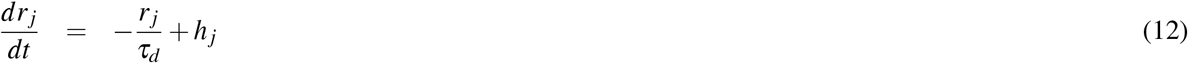

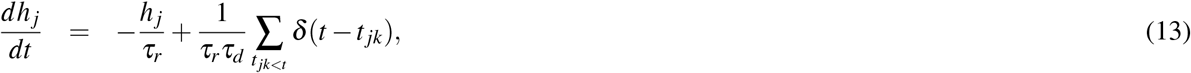

where *τ*_*r*_ = 2 ms is the synaptic rise time, *τ*_*d*_ = 20 ms is the synaptic decay time and *t* _*jk*_ is the time at which the neuron *j*th fired spike *k*th. For other synapse types, see^59^.

**Table 2.**
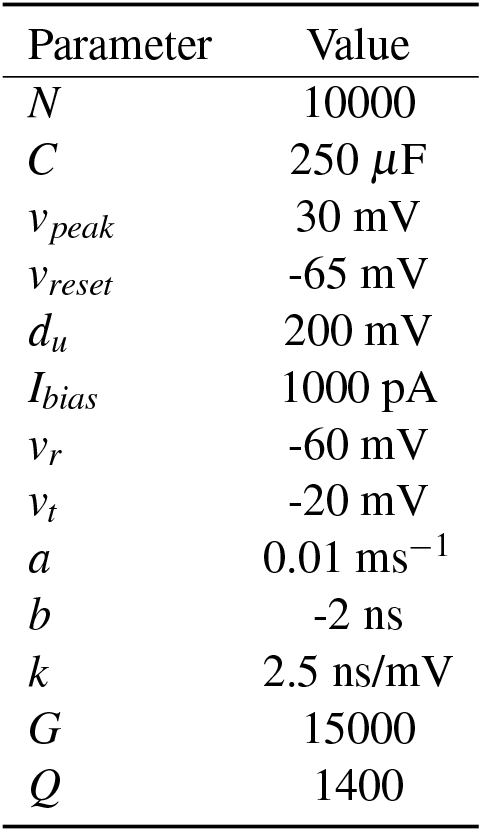
Neural parameters used to train the spiking neural network, described in equations (9), (10) and (16).

| Parameter | Value |
| --- | --- |
| $N$ | 10000 |
| $C$ | 250 $\mu\text{F}$ |
| $v_{peak}$ | 30 mV |
| $v_{reset}$ | -65 mV |
| $d_u$ | 200 mV |
| $I_{bias}$ | 1000 pA |
| $v_r$ | -60 mV |
| $v_t$ | -20 mV |
| $a$ | 0.01 $\text{ms}^{-1}$ |
| $b$ | -2 ns |
| $k$ | 2.5 ns/mV |
| $G$ | 15000 |
| $Q$ | 1400 |

The output of a spiking neural network is defined as:

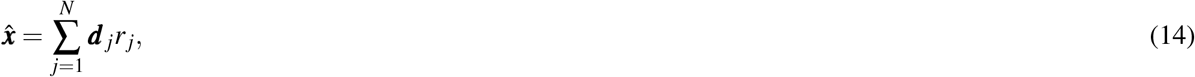

where ***d*** _*j*_ is an *m*-dimensional vector known as the linear decoder for the firing rate (see supplementary Figure 7D). Here, we want to train the network such that:

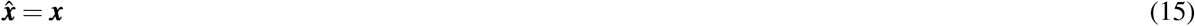

where ***x*** = (*x*_1_, *x*_2_, …, *x*_*m*_) are the desired dynamics or the supervisor that the network should mimic. Since the oscillators’ phases ***γ*** and ***ϕ*** are discontinuous and wrapped around the interval [0, 2*π*), the following supervisor for the chimera was used: ***x*** = (cos ***ϕ***, sin ***ϕ***, cos ***γ***, sin ***γ***). With 2*n* (*n* = 3) oscillators, this results in a *m* = 4*n* = 12 dimensional supervisor, see^61^ for details.

In order to achieve Eq. (15) we use the FORCE method^58^, which adds a second set of weights 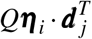 when defining the synaptic currents. Equation (11 can be rewritten as:

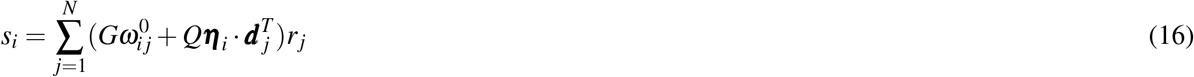

The FORCE method has three phases, the pre-learning, the learning and the post-learning. In the pre-learning phase, the initial synaptic connection matrix 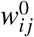 initializes the neurons’ dynamics into a well-known high-dimensional chaotic regime^79,80^. The matrix is static and sparse with each element drawn from a normal distribution with mean 0 and variance 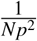 where *p* is the sparsity degree (set to 90% sparse or *p* = 0.1). The variable *G* controls the network’s chaotic behaviour and its value depends on the neuronal model, see^59^ for a detailed explanation. Here, we set *G* = 1.5× 10^3^.

The learning phase involves a second set of weights, given by 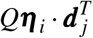. Where the parameter *Q* scales the encoding vector ***η***_*i*_, which has been drawn randomly and uniformly from [−1, 1]^*m*^ (where *m* is the dimensionality of the supervisor). By increasing *Q*, the feedback applied to the network is strengthened. A value of *Q* = 1.4 × 10^3^ was used for all simulations.

In the learning phase, the FORCE method enforces the aforementioned constrain 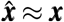 by changing ***d*** _*j*_ online (i.e., as the network is being simulated) with the Recursive Least Squares (RLS)^58^. RLS has an online solution for the optimal **d**, the one that minimizes the squared error **e** between the network output 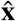 and the complex signal or supervisor **x**. RLS updates to **d** at each time step *n* are:

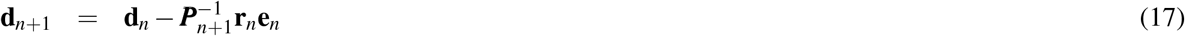

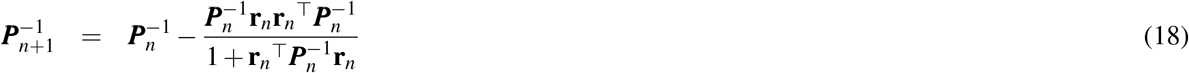

where **d**_0_ = 0 and ***P***_0_ = ***I***_*n*_*/λ*. The parameter *λ* controls the rate of the error^58^ and we set it to *λ* = 1. The parameter **I**_*n*_ is a *N* × *N* identity matrix

The third step of the FORCE method is the post-learning phase. RLS is turned-off and the weight matrix 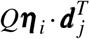 is no longer dynamic but static. The FORCE method is successful if the network is able to reproduce the supervisor for a fixed ***d***.

As a final step, we enforce Dale’s law in the spiking neuronal network. In Dale’s law, a neuron can only be either inhibitory or excitatory, not both. Dale’s Law was enforced by constraining ***ω*** to the inhibitory/excitatory nature of each individual neuron. If neuron *i* is inhibitory (excitatory), all of its outgoing connections will be negative (positive): 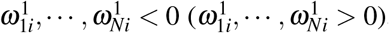, as seen in supplementary Figure 7A. We first define 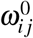such that 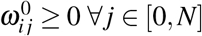 (the first half of the population of neurons only projects positive weights, i.e., excitatory neurons) and 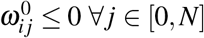 (the second half of the population of neurons only projects negative weights, i.e., inhibitory neurons). Second, the trained matrix 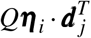 is limited to project either positive or negative weights. We obtain that by defining ***η*** as ***η*** = ***η***_−_ + ***η***_+_, where ***η***_−_ and ***η***_+_ are unequivocally defined as negative and positive matrices, respectively. And finally, ***d***^*T*^ is defined such that 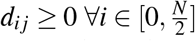 and 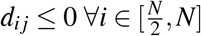 For the exact implementation refer to Additional Information where the link to the code is available and for more details see.^38, 59^

## Acknowledgements

W. N. was funded by a New Frontiers Research Foundation Exploration grant (NFRFE-2019-416 00159), an NSERC Discovery Grant, and a Hotchkiss Brain Institute start-up fund. J. D. was supported by the Natural Sciences and Engineering Research Council of Canada (RGPIN/05221-2020). M. M. thanks the Hotchkiss Brain Institute and the Cumming School of Medicine for their financial support.

## Author contributions statement

M. M, J. D., and W. N. conceived of the study. M. M performed all the numerical simulations. M. M., W. N., and J. D. wrote the manuscript.

## Additional information

There are no competing interests.

## Data availability

The datasets generated and analysed during the current study can be generated with the available codes. They are also available from the corresponding author on reasonable request.

## Code availability

Code is available from the corresponding author. No password required.

## Supplementary Material for

### Video 1

Representation of some nodes (we selected 20) of the chimera on a ring (500 in total). The node colour indicates the phase of each oscillator, which ranges from 0 to 2*π*. We observe what we would expect from the chimera definition, some are in sync and some are not. Available at: https://youtu.be/aY_ptuMoIkQ.

### Video 2

Same as Video 1, but for a different time. The chimera persists, but the synchronised sub-population drifts: some nodes change from a synchronised to an unsynchronised dynamics and vice versa. Available at: https://youtu.be/l0kB2qRgfUk.

**Supplementary Figure 1:**
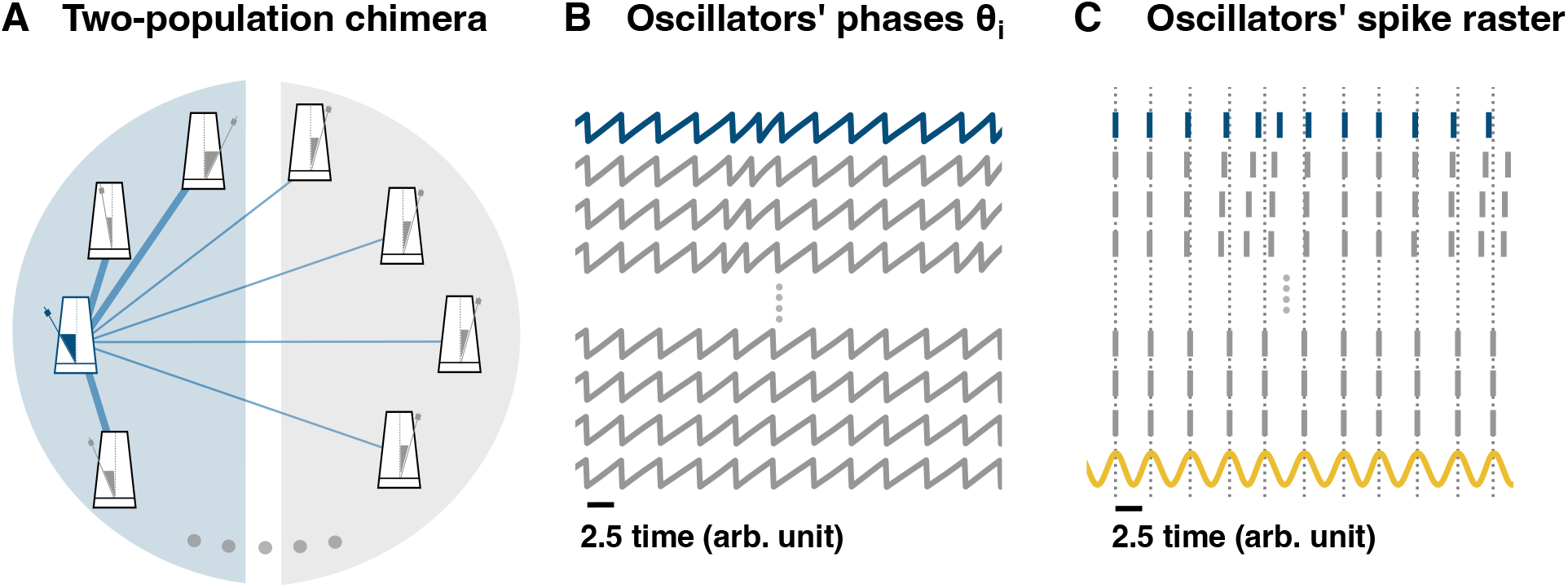
From a two-population chimera state to hippocampal phase precession. **A** Schematic representation of a two-population topology induced chimera. Non-local coupling is used, see main text for equations. For clarity, only the coupling for the blue oscillator (blue or dark grey metronome for b/w printing) has been depicted (blue or dark grey edges for b/w printing). Edge thickness represents the connection weight strength. **B** Time-series of different oscillators: some unsynchronized and some synchronized. **C** Oscillators’ spike raster plot: panel **B** transformed into a raster plot. See Methods (main text) for details. The sinusoidal curve (yellow) represents the theta oscillation. It is computed as cos(*ϕ*_*j*_), where *ϕ*_*j*_ corresponds to the phase from any of the synchronized oscillators. Dotted lines correspond to the peaks of the sinusoidal signal and to the spikes of the synchronized nodes.

**Supplementary Figure 2:**
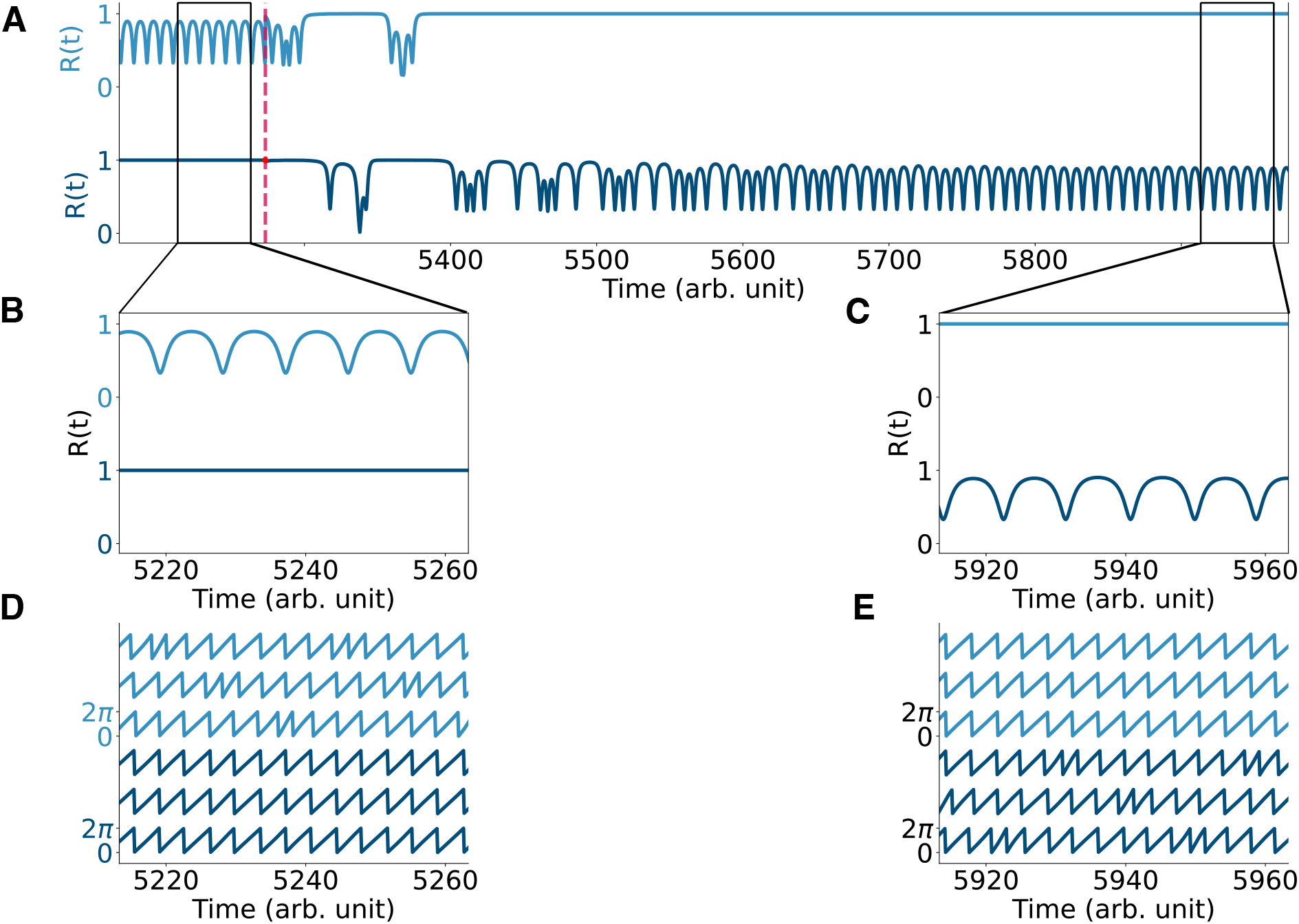
Perturbing the two-population chimera. **A** Order parameter R(t) for the two populations chimera ***ϕ*** (light blue, top) and ***γ*** (dark blue, bottom) before and after the system is being perturbed (dashed red line). The order parameter is computed at each time step as 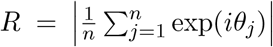 and it quantifies the synchronization of any oscillatory system with phases 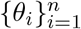. For synchronized systems |*R*| = 1 and for systems that are not fully synchronized, 0 ⩽ |*R*| *<* 1. **B, C** Zoom-in of the order parameter before and after the perturbation. **D, E** Time-series for the two populations chimera before and after the perturbation.

**Supplementary Figure 3:**
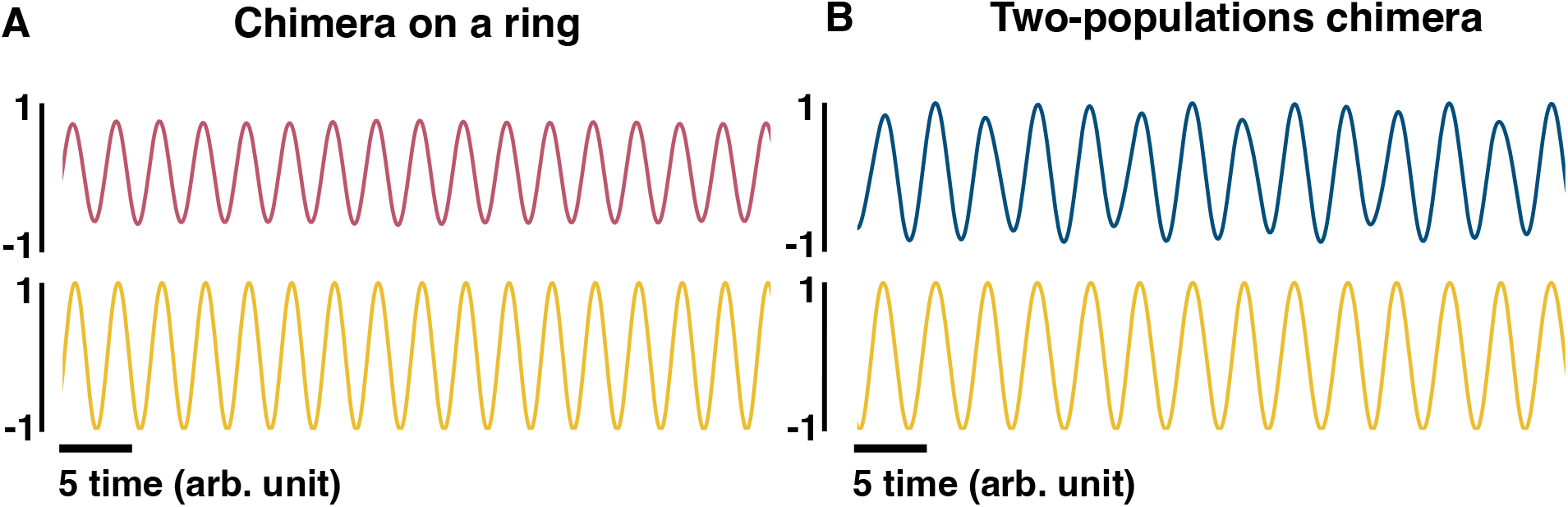
Phenomenological LFP. **A** The pink (top) sinusoidal curve is computed as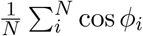, where *ϕ*_*i*_ is the phase of oscillator *i* and can be regarded as an equivalent to the lfp for the chimera on a ring. The yellow (bottom) sinusoidal curve is computed as cos *ϕ*_*s*_ where *ϕ*_*s*_ is the phase of one of the synchronized oscillators, and can be regarded as an LFP. **B** The blue (top) sinusoidal curve is computed as 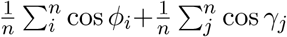 where *ϕ*_*i*_ and *γ*_*i*_ are the phases of oscillators *i* and *j*, respectively, and can be regarded as an equivalent LFP for the two-population chimera. The yellow (bottom) sinusoidal curve is computed as cos *ϕ*_*i*_, given that *ϕ*_*i*_ belongs to the synchronized population.

**Supplementary Figure 4:**
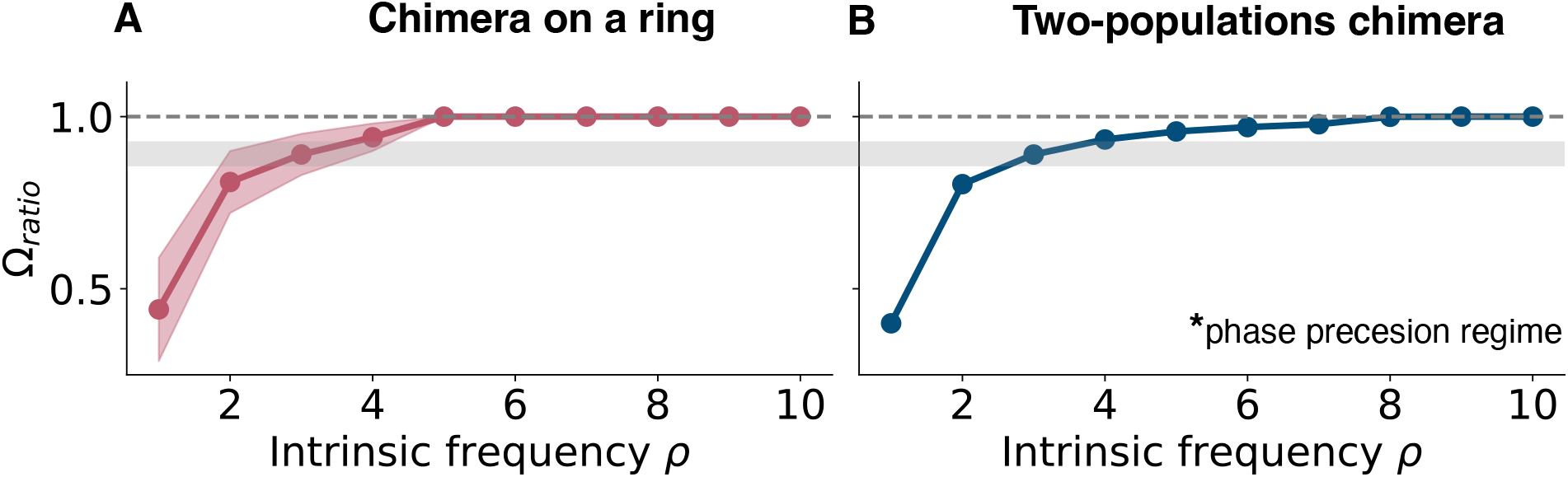
Mean phase velocity ratio for higher intrinsic frequencies. **A** Mean phase velocity ratio Ω_*ratio*_ for large values of the intrinsic frequency for the chimera on a ring. The pink region indicates the variation of the mean phase velocity ratio, since there isn’t a unique value for the mean phase velocity for the unsynchronized group. It is computed as the standard deviation of 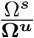. **B** Mean phase velocity ratio Ω_*ratio*_ for large values of the intrinsic frequency for the two populations chimera. Grey region on both panels: it indicates where both systems have their phase precession regime, i.e., Ω_*ratio*_ = 0.88.

**Supplementary Figure 5:**
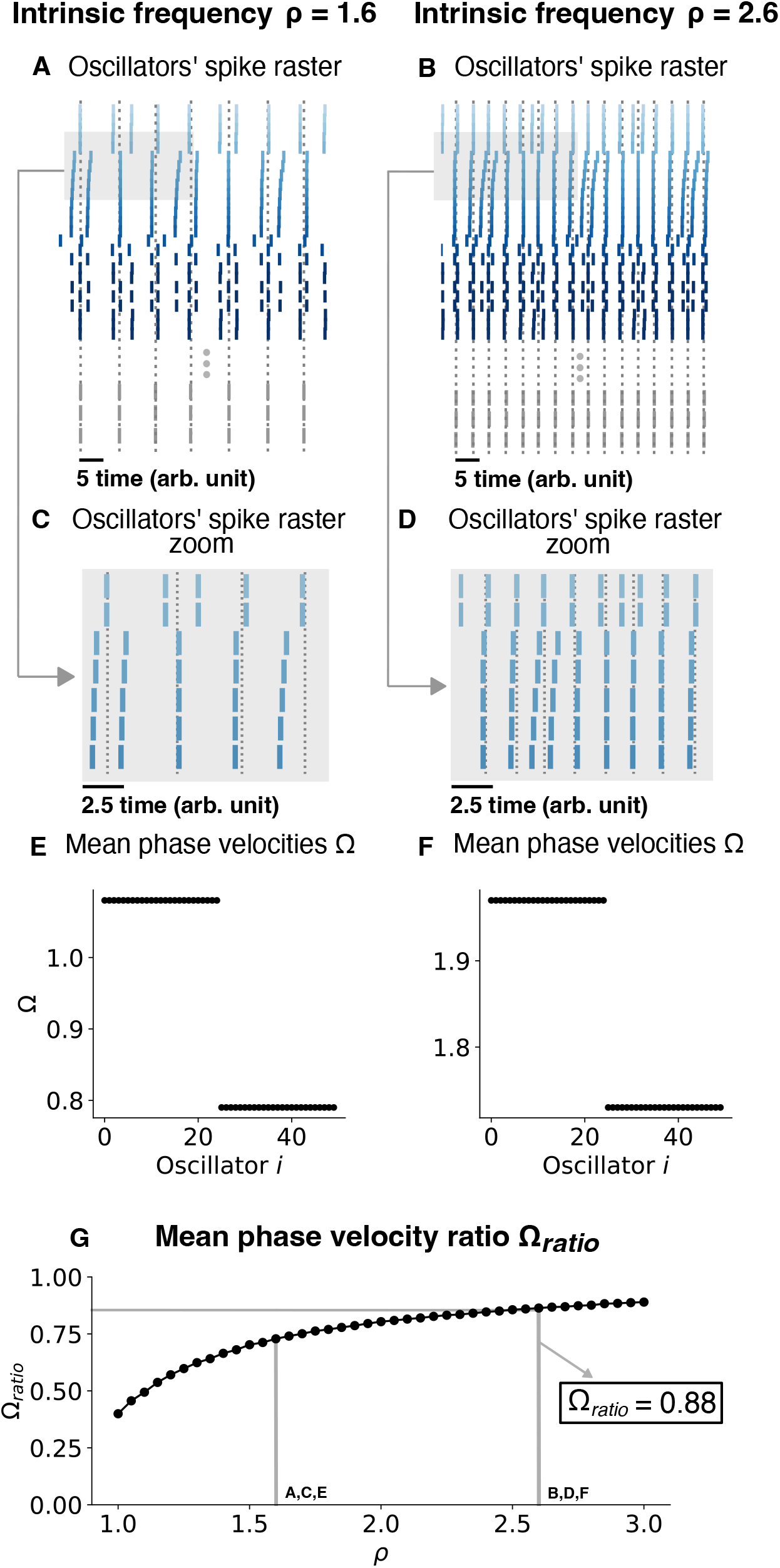
Changing the chimera state by changing the intrinsic frequency for a two-population chimera. **A, B** Oscillators’ spike raster plots for *ρ* = 1.8 and *ρ* = 2.8, respectively. Dotted lines correspond to the spikes of the synchronized nodes (grey ticks). **C, D** Respectively, zoom in on panels **A** and **B** (light grey box). **E, F** Mean phase velocity profile for *ρ* = 1.8 and *ρ* = 2.8, respectively. **G** Mean phase velocity ratio Ω_*ratio*_ in function of the intrinsic frequency *ρ*. Here *n* = 25 oscillators for each population.

**Supplementary Figure 6:**
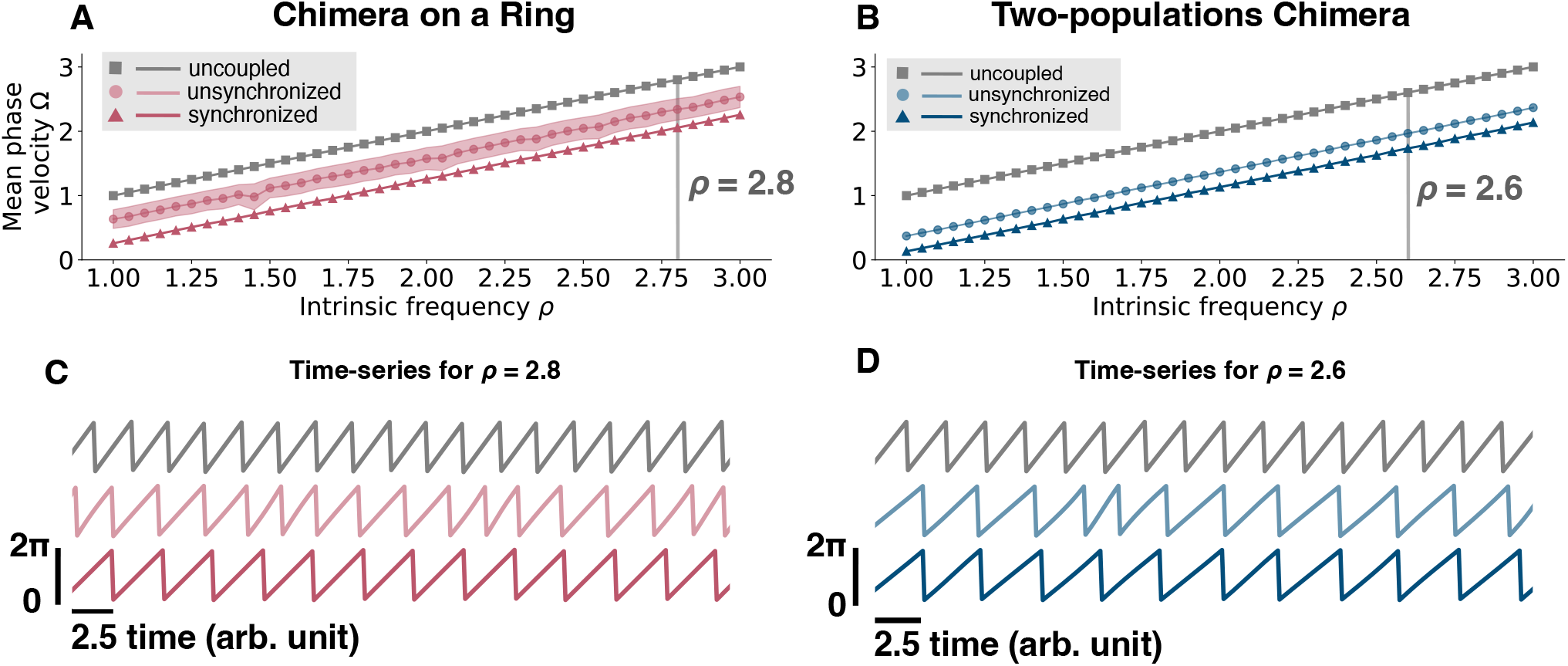
Mean phase velocity for an uncoupled oscillator and for different intrinsic frequencies. **A** Mean phase velocity in function of the intrinsic frequency *ρ* for an uncoupled oscillator, i.e. 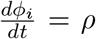(grey squares), for the unsynchronized oscillators (light pink circles), and for the synchronized oscillators (pink triangles) of the chimera on a ring. For the unsynchronized oscillators the mean phase velocity is computed as the mean of **Ω**^*u*^, since we get a different Ω^*u*^ for each oscillator *i*. The light pink region is computed as the standard deviation of **Ω**^*u*^. **B** Mean phase velocity in function of the intrinsic frequency *ρ* for an uncoupled oscillator, i.e. 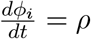 (grey squares), for the unsynchronized population (light blue circles), and for the synchronized population (blue triangles) of the two-populations chimera. **C** Time-series for *ρ* = 2.8 for the three different cases, uncoupled (grey, top), unsynchronized (light pink, middle) and synchronized (pink, bottom) for the chimera on a ring. **D** Time-series for *ρ* = 2.8 for the three different cases, uncoupled (grey, top), unsynchronized (light blue, middle) and synchronized (blue, bottom) for the two-populations chimera.

**Supplementary Figure 7:**
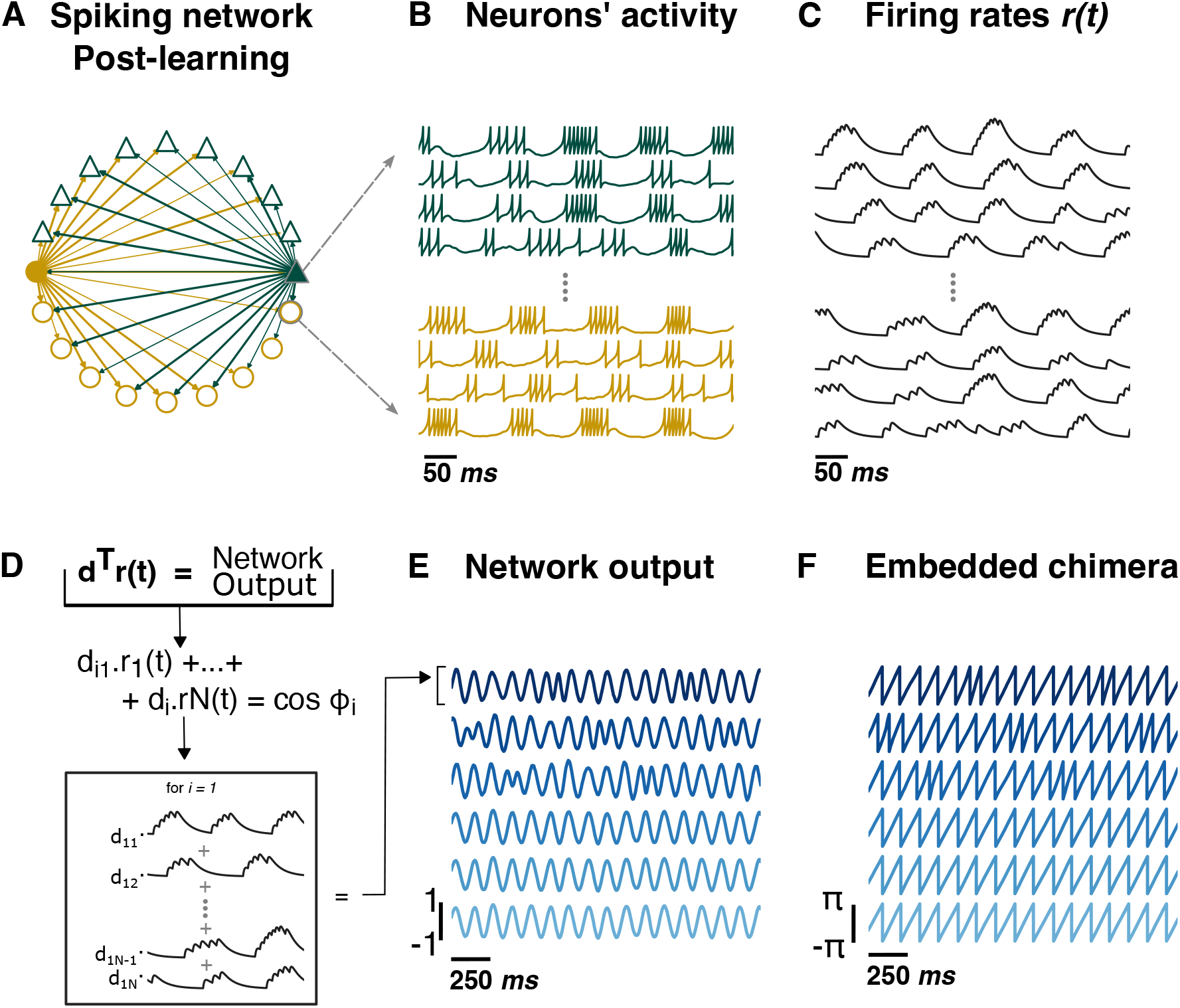
Supplementary Figure 4: Training a spiking neural network to output a chimera state. **A** Spiking neural network. Each node represents either an excitatory (green or dark grey triangle for b/w printing) or inhibitory (yellow or light grey circle for b/w printing) neuron. For clarity, only the connections for two neurons (filled triangle and circle) are depicted. The network respects Dale’s law: an excitatory (inhibitory) neuron will only excite (inhibit) its connections, regardless of the neuron target type. As a result, excitatory (inhibitory) neurons just have green or dark grey for b/w printing (yellow or light grey for b/w printing) outgoing connections. **B** Voltage traces for excitatory (green or dark grey triangle for b/w printing) and inhibitory (yellow or light grey round for b/w printing) neurons. **C** Firing rates ***r*(*t*)** obtained from filtering the spikes with a two double exponential filter, see equations for details. **D** The network output is given by **d**^*⊤*^***r*(*t*)**, which is a *s* x *n*_*t*_ matrix (*s* is the total number of supervisors). Each network output column *i* is *n* time units long and is a linear combination of the firing rates *r*_1_(*t*),, *r* (*t*) with *d*_iN_ as coefficients. **F** Network output 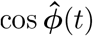 and cos 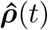. **F** Embedded Chimera from network output: 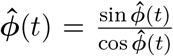 and 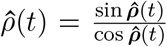. We recover the two-populations chimera (where we had *n* = 3 oscillators per population).

## Notes

### Competing Interest Statement

The authors have declared no competing interest.

